# Language network connectivity increases in prodromal Alzheimer’s disease

**DOI:** 10.1101/2020.11.22.393199

**Authors:** A. Pistono, M. Senoussi, L. Guerrier, M. Rafiq, M. Gimeno, P. Péran, M. Jucla, J. Pariente

## Abstract

Language production deficits occur early in the course of Alzheimer’s disease (AD); however, only few studies have focused on language functional networks in prodromal AD. The current study aims to uncover the extent of language alteration at a prodromal stage, on a behavioral, structural and functional level, using univariate and multivariate analyses. Twenty-four AD participants and 24 matched healthy controls underwent a comprehensive language evaluation, a structural T1-3D MRI and resting-state fMRI. We performed seed-based analyses, using the left inferior frontal gyrus and left posterior temporal gyrus as seeds. Then, we analyzed connectivity between executive control networks and language network in each group. Finally, we used multivariate pattern analyses to test whether the two groups could be distinguished based on the pattern of atrophy within the language network; atrophy within the executive control networks, as well as the pattern of functional connectivity within the language network; and functional connectivity within executive control networks. AD participants had language impairment during standardized language tasks and connected-speech production. Univariate analyses were not able to discriminate participants at this stage, while multivariate pattern analyses could significantly predict the group membership of prodromal patients and healthy controls, both when classifying atrophy patterns or connectivity patterns of the language network. Language functional networks could discriminate AD participants better than executive control networks. Most notably, they revealed an increased connectivity at a prodromal stage. Multivariate analyses represent a useful tool for investigating the functional and structural (re-)organization of the neural bases of language.

**Highlights:** Language network connectivity discriminates prodromal AD from healthy controls

Language network connectivity increases in prodromal AD

Atrophy patterns in the language network do not correlate with connectivity patterns in AD

## 1. Introduction

Language production deficits occur early in the course of Alzheimer’s disease (AD). Most studies have shown impairment in fluency tasks and confrontation naming tasks (Taler and Phillips, 2008), usually attributed to lexical-semantic impairment (Joubert et al., 2010). These tasks have also been shown to accurately discriminate prodromal patients from healthy controls (Mueller et al., 2016; Taler and Phillips, 2008). Fewer studies have analyzed other language processes. Some studies have shown preserved syntactic abilities in early AD (Taler and Phillips, 2008), while others did not find such preservations (Kemper et al., 1993). Most studies have stressed the fact that phonological capacities are relatively preserved in early AD (Taler and Phillips, 2008). More and more studies have been focusing on connected speech production in AD, for the assessment of the functional use of language and cognition. They revealed several impairments in AD: reduced lexical content (Ahmed et al., 2013), increased word-finding difficulty and use of repetitions and self-corrections (de Lira et al., 2011), etc. While most studies focused on AD at a dementia stage, other studies revealed changes as early as the prodromal stage. Mueller et al. (2016) demonstrated that prodromal patients had lower lexical richness compared to healthy controls, but similar production of filled pauses (e.g. “*hm*”). Pistono et al. (2018) also showed that these patients produced more modalizing discourse, which refers to “discourse about discourse” (i.e. comments, feelings and uncertainty about the task).

Neuroimaging studies in AD patients have shown that language impairments are associated with atrophy or hypometabolism in the left inferior frontal gyrus (IFG) and temporal regions (e.g. Melrose et al., 2009). However, besides the alteration of isolated brain regions, the functional connectivity within brain networks can underlie the cognitive impairments or compensations observed. Resting-state functional connectivity is one of the current methods that allows functional brain networks to be investigated, including language functional network (e.g. Muller et al., 2016; Muller & Meyer, 2014). In AD, only few studies focused on this network, reporting lower functional connectivity in AD (Weiler et al., 2014, patients’ mean MMSE: 18.86; Mascali et al., 2018, patients’ mean MMSE: 20.5) and prodromal AD (Montembeault et al., 2019, patients’ mean MMSE: 24.9) compared to healthy controls. These studies used the left IFG (Mascali et al., 2018; Montembeault et al., 2019) or left posterior temporal gyrus (Mascali et al., 2018; Montembeault et al., 2019; Weiler et al., 2014) as a seed. They also showed that connectivity changes were only marginally correlated with AD participants’ language performance (i.e. no significant correlations in Mascali et al., 2018, no correlations with IFG’s connectivity map in Montembeault et al., 2019). However, it is possible that some changes remain unnoticed when focusing exclusively on the language network. For example, we now know that, in healthy aging, the language network interacts with the executive control network/attentional network to maintain a sufficient level of language performance (Hoffman & Morcom, 2018; Pistono et al., 2020). It is therefore possible that prodromal AD is primarily characterized by a loss of this compensation, rather than an alteration within the language network.

Second, univariate fMRI analyses may not be able to uncover the extent of changes occurring at a prodromal stage. Indeed, analysis of structural or functional MRI data is traditionally performed in a univariate manner, where each voxel or area in the brain is separately tested for a condition of interest. By contrast, multivariate pattern analyses (MVPA) simultaneously consider patterns of information (i.e. atrophy or BOLD signal), leveraging the multivariate, i.e. multi-voxel, and distributed nature of neural representations (Haynes and Rees, 2006). In other words, while univariate analyses ask to what degree each voxel’s activity is affected by a particular condition, MVPA examines whether, by contrast, an experimental manipulation or a clinical population can be predicted based on the pattern of activity across a set of voxels. Using multivariate patterns of activity, i.e. activity across multiple voxels, can increase sensitivity in differentiating between individuals or conditions (Haynes & Rees, 2006; but see Hebart & Baker (2018) for a discussion on the benefits and pitfalls of MVPA as compared with classical univariate analyses). Regarding Alzheimer’s disease, Liu et al., (2018) applied MVPA to investigate the topologic alterations of resting-state functional connectivity in participants with subjective cognitive decline, prodromal AD and AD compared with healthy individuals. They showed that by using MVPA, it was possible to predict whether a participant belonged to one of the three clinical groups or to the healthy control group, which indicated that patterns of resting-state data are already discriminant for cognitive decline and prodromal AD. Further work is required to understand how these changes relate to patients’ cognitive impairment.

In the current study, we focus on language processing to uncover the extent of language alteration at a prodromal stage on a behavioral, structural and functional level, using univariate and multivariate analyses. Additionally, we will examine whether structural and functional changes are correlated with language performance, using both standardized and connected speech tasks. Regarding language performance, we expect behavioral inter-group differences for both the standardized language tasks and discourse task, in line with current literature on prodromal AD (e.g. Mueller et al., 2016). Regarding functional connectivity, we will first analyze language networks using the same method as previous literature on AD, and the same two seeds: left IFG and left posterior temporal gyrus (e.g. Mascali et al., 2018). We anticipate marginal inter-group differences with this analysis. We will then analyze connectivity between executive control networks and language network. We expect lower between-network connectivity in prodromal AD participants, correlated with lower language performance. Finally, we will use MVPA to test whether it is possible to distinguish the two groups based on (i) the pattern of atrophy within the language network, (ii) atrophy within the executive control networks, as well as (iii) the pattern of functional connectivity within the language network and (iv) functional connectivity within executive control networks (using atlases from Shirer et al., 2012). Based on previous studies showing that functional connectivity is affected in prodromal AD, we predict that both structural and functional information will allow to discriminate AD participants from healthy controls using MVPA. We also hypothesize that functional changes within both language and executive control networks will be related to language performance. In particular, lower lexical performance (i.e. naming and fluency tasks) and lexical content during connected speech production will be correlated with functional connectivity alteration in the AD group.

## 2. Material and Methods

### 2.1. Participants

Participants were right-handed and native French speakers with no history of neurological or psychiatric problems. In order to avoid possible reorganization of the language network due to multilingualism, we only included speakers that did not have a good command and/or a frequent use of a language other than French. All the participants provided written, informed consent before participating in the study and received monetary compensation for their participation. The current study was approved by the ethics committee (IDRCB: 2015-A01416-43).

AD participants were selected if they presented with a memory complaint and had no concomitant history of neurological or psychiatric disease. They underwent the following pre-inclusion assessment:

– Autonomy in daily living (Instrumental Activities of Daily Living (IADL), Graf, 2008);
– Global cognition (Mini-Mental State Evaluation (MMSE));
– Anterograde verbal memory (Free and Cued Selective Reminding Test (FCSRT, Van der Linden et al., 2004)).
– Amyloid assessment with cerebrospinal fluid (CSF) analysis by lumbar puncture: CSF biomarker levels of total tau (T-Tau), phospho-tau (P-Tau), Ab42 and Ab40 were measured using an ELISA method (Innogenetics, Ghent, Belgium). Innotest Amyloid Tau Index (IATI) was calculated. P-Tau ≥ 60 pg/ml and IATI ≤ 0.8 were deemed to be suggestive of AD. In case of an ambiguous profile (P-Tau < 60 pg/ml or IATI > 0.8), we calculated the Ab42/Ab40 ratio and a score < 0.045 was considered to be compatible with a diagnosis of AD.

Individuals with AD were included if they met the following criteria: MMSE ≥ 24; IADL < 1 and based on the IWG-2 criteria (Dubois et al., 2014): evidence of a gradual and progressive change in memory function reported by patient or informant for more than 6 months and demonstrated by an episodic memory test, and CSF evidence of AD.

Matched healthy control participants underwent the same pre-inclusion neuropsychological assessment as the AD group. They were included if they had no memory complaint and no history of neurological or psychiatric disease and a MMSE ≥ 27. They were excluded if they presented with cognitive impairment (test scores < -1.5 SDs) during the pre- or post-inclusion neuropsychological assessment.

### 2.2. Cognitive evaluation

#### 2.2.1. Neuropsychological assessment

All participants also underwent a comprehensive neuropsychological assessment. Visual recognition memory was assessed with the Doors and People test (Baddeley et al., 1994). Short-term memory and working memory were evaluated with the WAIS-III Digit Span and Backward Digit Span subtest (Wechsler, 1997). Cognitive flexibility was assessed with the Trail Making Test (TMT, Reitan, 1958). Praxis was explored with Mahieux’s assessment (Mahieux-Laurent et al., 2008) and gnosis with the Visual Gnosis Evaluation Protocol (VGEP, Agniel, Joanette, Doyon, & Duchein, 1992). Apathy and depression were also measured, using the Starkstein scale (Starkstein et al., 1992) and the Beck Depression Inventory (Beck et al., 1961).

#### 2.2.2. Language assessment

Language was assessed with the GREMOTs assessment (Bézy et al., 2016). GREMOTs is a computerized battery of language tests that evaluates both oral and written language as well as production and comprehension at different levels (i.e. phonological processing, lexical processing and syntactic processing).

This battery includes a connected-speech task, which we analyzed more specifically. With regards to the procedure for this task, the participants were given the same instructions: “This is a story depicted in 5 pictures. Tell me the story with as many details as possible.” During the task, the experimenter remained neutral and avoided speaking in order to ensure uniform conditions for discourse production. The oral productions of participants were recorded and manually and orthographically transcribed. The following variables were used to analyze the discourse of both the AD group and the cognitively normal controls:

– Total number of words in the narrative;
– Lexical content: proportion of closed class and open class words (i.e. nouns, most verbs, adjectives, numerals and adverbs of manner). Standardized indexes were calculated according to the following formula: (Open class – Closed class)/(Open class + Closed class), similarly to Pistono et al., (2019);
– Proportion of self-corrections: number of self-corrections normalized per 100 words (e.g. when the speaker stops and resumes with a substitution for a word or a new utterance);
– Proportion of repetitions: number of repetitions (of sounds, syllables, words or partial phrases) normalized per 100 words;
– Proportion of filled pauses: number of filled pauses (e.g. “*hm*,” “*um*,” “*pff*”) normalized per 100 words;
– Proportion of modalizing discourse: number of words that are part of a modalizing utterance, normalized per 100 words (e.g. “*It seems that*”; “*I don’t know how to say it*”; etc.).

Intergroup comparisons for the neuropsychological assessment and the language assessment were performed using Student’s t-test for independent samples. Bonferroni-Holm corrections for multiple comparisons were applied.

### 2.3. Structural and functional MRI

#### 2.3.1. MRI Acquisition

MRI scans were performed for all participants using a 3-T imager (Philips Achieva dStream, Inserm/UPS UMR1214 ToNIC Technical Platform, Toulouse, France). A 3D-T1 image was acquired for anatomical reference with the following parameters: TR = 8 ms, TE = 3.7 ms, flip angle = 8°, matrix size = 256 x 256 mm, 170 slices, voxel size= 0.9 mm x 0.9 mm x 1 mm, slice thickness = 1 mm. Whole-brain resting-state fMRI images were obtained with the following parameters: TR = 2837 ms, TE = 40 ms, flip angle = 90°, 46 interleaved acquisition, slice thickness = 3 mm, matrix size = 80 x 80 mm, 200 volumes, total scan time 10 min. During scanning, participants were instructed to keep their eyes closed but to stay awake and avoid thinking of anything in particular. All participants affirmed that they were fully awake during the 10 minutes of the scanning.

#### 2.3.2. Preprocessing

The data were analyzed using the Conn toolbox (Version 18b, Whitfield-Gabrieli & Nieto-Castanon, 2012), implemented in MATLAB. The preprocessing pipeline of the functional images included: functional realignment and unwarp, slice-timing correction, outlier identification, normalization to the MNI template, and smoothing with a Gaussian kernel of 6 mm. This step created a scrubbing covariate (containing the potential outliers scans for each participant) and a realignment covariate (containing the six head motion parameters). Average realignment (*t*(46)=0.97, *p=0*.*17*) and maximum realignment (*t*(46)=1.16, *p=0*.*13*) did not significantly differ between the two groups. Then, the six head motion parameters plus their associated first-order derivatives, the identified outliers scans, white matter and cerebrospinal fluid signals and the effect of rest were removed by means of the CompCor method. The resulting preprocessed images were band-pass filtered (0.01 Hz–0.1 Hz) to remove physiological high-frequency noise (e.g. cardiac and respiratory fluctuations). Atlases were then masked with the participant’s gray matter mask. With this method, each ROI was restricted to voxels belonging to an estimated gray matter mask derived from the T1 segmentation.

#### 2.3.3. Voxel based morphometry (VBM)

Gray matter density was assessed using a voxel-based morphometry method on Statistical Parametric Mapping version 12 (SPM 12, Wellcome Trust Centre for Neuroimaging). For each participant, the 3D-T1 sequence was segmented to isolate gray matter and white matter partitions, modulated for deformation, normalized to the MNI space and smoothed (8×8×8 mm). Inter-group comparisons were then performed (voxel level p<0.05, FWE-corrected, cluster = 50 voxels).

#### 2.3.4. Seed-based analyses

The left Inferior frontal gyrus (LIFG) and the left posterior temporal gyrus (LSTG, including parts of the left middle/superior/supramarginal gyrus) were used as seeds, based on Shirer’s functional atlas of language (Shirer et al., 2012). Correlation maps were constructed by correlating the average BOLD-signal dynamic of the region of interest with the BOLD-signal of every other single voxel. To enforce a Gaussian distribution of the correlation data, Pearson’s correlation coefficients were then transformed to z-scores using the Fisher r to z transformation for subsequent t-tests. These individual z values maps were entered into a one-sample t-test to determine the functional network correlated with spontaneous activity of the seed region within each group (p < 0.05 FWE at the cluster level). We then performed two-sample t-tests to detect inter-group differences. The threshold for second-level maps was set at p < 0.05 FWE at the cluster level.

#### 2.3.5. Within- and between-network connectivity

To measure within-network and between-networks connectivity, we selected networks from Shirer’s atlas (2012): language network, left executive control network (left ECN), right executive control network (right ECN).

The language network includes 7 ROIs within the left IFG, right IFG, left middle temporal gyrus, left middle/angular gyrus, left middle/superior/supramarginal gyrus, right middle/superior/ supramarginal gyrus, left thalamus and left cerebellum. The left ECN includes 6 ROIs within the left middle frontal/superior frontal gyrus, left IFG/orbitofrontal gyrus, left superior/inferior parietal/precuneus/angular gyrus, right cerebellum and left thalamus. The right ECN includes 6 ROIs as well: the right middle frontal/superior frontal gyrus, right middle frontal gyrus, right superior frontal gyrus, right inferior parietal/supramarginal/angular gyrus, left cerebellum and right caudate.

Within- and between-network connectivity (average for all the ROIs within each network) was evaluated for each participant. More precisely, within-network connectivity is a mean composite network connectivity estimate, calculated by means of pairwise correlations between all the regions comprising an individual network. Between-network connectivity is the result of pairwise correlations between the regions in each pair of different networks. Averages of within- and between-network connectivity were compared between groups with one-tailed t-tests to assess whether healthy controls present greater within- and between-network connectivity than AD participants.

#### 2.3.6. Multivariate pattern analyses

To investigate whether the two groups could be identified based on the pattern of atrophy or connectivity within the language network and the ECN networks, we performed multivariate pattern classification.

Supervised classification analyses, performed using a classifier algorithm, consist in training a classifier to distinguish two or more classes of data (e.g. class 1: healthy controls (HC), class 2: patients (AD)) from a set of training samples by providing the corresponding labels of each sample, e.g. “healthy control” or “patient.” Following this training phase, the classifier is then tested on a test dataset composed of samples not used during the training phase, in order to assess whether the classifier is able to generalize to new unseen data. If the classifier is able to predict the class of novel samples in the test dataset, i.e. accurate prediction, it indicates that the multivariate pattern of information is informative about the classes of interest. To ensure unbiased evaluation of classification performance, this procedure is repeated over multiple independent divisions of the entire dataset into training and test datasets, i.e. cross-validation. The accuracy of classifier predictions, i.e. 0 for incorrect and 1 for correct, are then averaged across cross-validation folds to obtain a classification score between 0 and 1 (or 0% and 100%) that can be compared to chance level. For our analyses, there was always 2 classes, corresponding to the healthy control or patient groups, therefore chance level was 1/2 = 50%.

##### Features selection

Classifiers are sensitive to the ratio between the number of variables, e.g. voxels, and number of samples, i.e. the different data samples provided, which can cause overfitting and/or poor classification accuracy (Pereira et al., 2009). One method to prevent this is to perform the analysis on specific ROIs based on anatomical or functional data (Pereira et al., 2009). Doing so decreases the number of voxels used by the classifier and focuses on appropriate regions that allow for best discrimination. We therefore extracted ROIs from the Shirer’s atlas (Shirer et al., 2012) of language network (7 ROIs), left ECN (6 ROIs) and right ECN (6 ROIs) to perform 6 classifications: gray matter density within areas of each of these three networks, as well as functional connectivity between areas of each of these three networks. To extract each participant’s gray matter density within each ROI, we performed a one sample t-test (using SPM12) using each network as an inclusive mask. To extract individual connectivity values between each ROI of the networks under study, we performed a one sample t-test within each network, using the Conn toolbox.

##### Classification procedure

We used a linear discriminant analysis (LDA) classifier implemented in the Scikit-learn toolbox (Pedregosa et al., 2011). More precisely, we trained the LDA classifier to distinguish the two classes of data, i.e. “healthy controls” versus “patients.” The classification was performed in a leave-one-out cross-validation approach. In each cross-validation fold, the classifier was trained on data from all but one participant and used on the left-out participant to predict its class membership. This procedure was repeated until each trial’s class had been used as a test.

##### Permutation test

To evaluate the significance of classification accuracies, for each analysis, we computed permutation tests. In order to estimate the null distribution of classification accuracy, we randomly permuted the labels of all samples (i.e. HC or AD) and performed the classification analysis 100,000 times, yielding 100,000 surrogate classification accuracies under the null hypothesis that the two classes are completely interchangeable. From these surrogate distributions, we computed the probability if observing a certain classification accuracy, i.e. p-value.

##### Feature contribution

For each classification, we extracted each feature contribution by using a method that allows an “informativity” measure to be extracted from classifier weights (Haufe et al., 2014). Indeed, classifier weights cannot be interpreted, as they reflect both noise and signal in the data; we thus used this approach to evaluate the extent to which a certain feature was informative in performing the classification. For each classification, the contribution value of each feature was calculated. Furthermore, a null distribution of each feature’s contribution was computed using the permutation procedure described above to estimate the significance of the contribution values.

#### 2.3.7. Correlations between functional connectivity and language performance

For the different functional analyses (i.e. seed-based, between-network connectivity, MVPA), significant inter-group differences were further examined through intra-group correlations. To do so, we chose the most sensitive variables during the language assessment (object naming, famous face naming, word spelling, written semantic verification, sentence spelling and text comprehension) and the narrative task (lexical content, modalizing discourse, self-corrections). We performed Kendall correlations and then applied Bonferroni-Holm corrections.

## 3. Results

### 3.1. Population

Twenty-four AD participants and 24 healthy controls were recruited. Both groups were matched for age (AD group: 72.9±8 years old; HC group: 70±4 years old, p=0.09), gender (AD group: 13 women; HC group: 11 women) and level of education (years of education, AD group: 12.5±4; HC group: 12.4±4, p=0.9).

During the pre-inclusion assessment, patients had a lower MMSE (AD group: 25.5±2.6; HC group: 29±1, p<0.0001) and lower performance during the FCSRT than the control group (sum three free recalls AD group: 14.17±9.69; HC group: 32.29±4.79, p<0.0001; sum three cued recalls AD group: 30.33±12; HC group: 46.42±1.93, p<0.0001).

#### 3.1.1. Neuropsychological assessment

During the post-inclusion assessment, AD participants’ performance on the Doors and People test, digit span forward and Trail Making Test was also lower than that of the control group, as shown in Table 1.

**Table 1.** Performance during the neuropsychological assessment. Results represent mean±SD. Results that are significant after Bonferroni-Holm correction are in bold. Cohen’s d values were measured for these variables only.

|  | Healthy<br>Controls | AD<br>participants | p value | Cohen's<br>d |
| --- | --- | --- | --- | --- |
| **Doors and People test, set A** | $10.78 \pm 1.38$ | $8.00 \pm 2.55$ | **<0.0001** | **1.36** |
| **Digit span forward** | $6.00 \pm 1.00$ | $5.21 \pm 0.98$ | **0.009** | **0.80** |
| **Digit span backward** | $4.83 \pm 1.40$ | $4.04 \pm 0.91$ | 0.027 | - |
| **Trail Making Test, A** | $38.79 \pm 12.50$ | $51 \pm 16.41$ | **0.006** | **0.83** |
| **Trail Making Test, B-A** | $55.13 \pm 27.86$ | $114.22 \pm 81.83$ | **0.002** | **0.97** |
| **VGEP** | $35.26 \pm 1.10$ | $33.79 \pm 2.87$ | 0.026 | - |
| **Beck** | $2.58 \pm 2.21$ | $3.29 \pm 3.28$ | 0.384 | - |
| **Starkstein** | 9.50±4.19 | 11.78±4.60 | 0.082 | - |

#### 3.1.2. Brain atrophy

AD participants had significant atrophy in two clusters compared to the control group (see Appendix), with one cluster encompassing the left hippocampus, parahippocampus and thalamus (K voxels=3278; t=7.28; pFWE-corr<0.0), and one cluster involving the contralateral areas (K voxels=1076; t=7.40; pFWE-corr<0.05).

### 3.2. Language evaluation

#### 3.2.1. Standardized assessment

AD participants had lower performance during several lexical tasks, as well as syntactic tasks. Results are detailed in Table 2.

**Table 2.** Performance during the language assessment. Results represent mean±SD. Results that are significant after Bonferroni-Holm correction are in bold. Cohen’s d values were measured for these variables only.

|  |  | **Healthy**<br>**Controls** | **AD**<br>**participants** | **p value** | **Cohen's**<br>**d** |
| --- | --- | --- | --- | --- | --- |
| **Lexical**<br>**processing** | Repetition, words (/10) | 9.38±1.01 | 9.17±1.05 | 0.488 | - |
|  | Grammatical fluency<br>(category: verbs) | 35.21±11.66 | 27.08±11.23 | 0.018 | - |
|  | Semantic fluency<br>(category: fruits) | 19.33±3.38 | 15.04±6.03 | 0.004 | - |
|  | Phonemic fluency<br>(letter V) | 17.29±6.12 | 17±8.01 | 0.888 | - |
|  | Object naming (/36) | 34.7±1.40 | 32.63±1.91 | **<0.0001** | **1.23** |
|  | Action naming (/36) | 33.13±3.25 | 31.13±2.8 | 0.028 | - |
|  | Famous face naming<br>(/10) | 8.75±1.15 | 4.83±2.78 | **<0.0001** | **1.84** |
|  | Reading, words (/30) | 29.71±.55 | 29.33±.92 | 0.092 | - |
|  | Spelling, words (/12) | 11.58±.504 | 10.04±1.33 | **<0.001** | **1.53** |
|  | Oral semantic verification (/18) | 17.04±1.27 | 15.96±1.6Å | 0.013 | - |
|  | Written semantic verification (/18) | 16.3±1.69 | 14±2.21 | **<0.001** | **1.17** |
| **Syntactic processing** | Repetition, sentences (/4) | 3.46±.78 | 3.42±.65 | 0.842 | - |
|  | Order execution (/6) | 5.96±.20 | 5.79±.42 | 0.084 | - |
|  | Sentence production (/6) | 5.75±.68 | 5.25±.94 | 0.040 | - |
|  | Syntactic comprehension (/24) | 21.25±2.51 | 18.92±3.62 | 0.013 | - |
|  | Spelling, sentence (/27) | 25.83±1.05 | 24.25±1.98 | **0.001** | **1.00** |
|  | Text comprehension (time in seconds) | 49.3±15.73 | 80.88±30.51 | **<0.0001** | **1.30** |
| **Phonological processing** | Repetitions, non-words (/6) | 5.54±.66 | 5.08±.93 | 0.055 | - |
|  | Reading, non-words (/15) | 14.67±.64 | 13.79±1.06 | **0.001** | **1.00** |
|  | Spelling, non-words (/6) | 5.50±.59 | 4.96±1.04 | 0.032 | - |

#### 3.2.2. Connected-speech production

The AD group did not produce shorter narratives compared to healthy controls (number of words AD group: 172±112; HC group: 146±84, p=0.4). However, they produced significantly more self-corrections (AD group: 3.3±2.1; HC group: 1.7±1.5, p=0.006, Figure 1) and more modalizing discourse (AD group: 12.1±11.5; HC group: 3.7±7.5, p=0.005) while performing this task. Their lexical content was also lower than the control group (AD group: -0.82±0.8; HC group: 0.19±0.8, p=0.0001, Figure 1). On the contrary, the two groups produced the same proportion of repetitions (AD group: 1.9±1.9; HC group: 1.2±0.9, p=0.1) and filled pauses (AD group: 4.1±2.8; HC group: 3.5±2.9, p=0.5).

**Figure 1.**
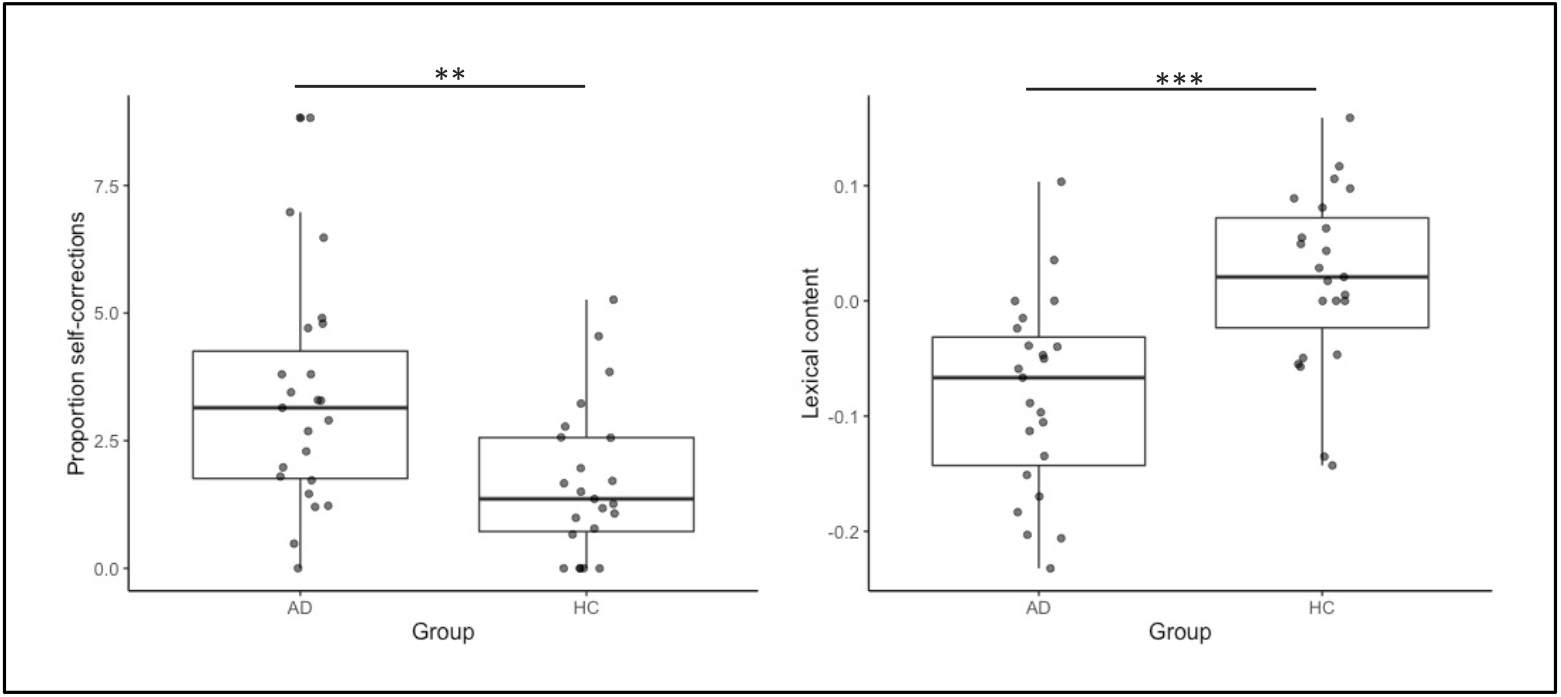
Inter-group comparisons for self-corrections (left) and lexical content (right) between AD participants (AD) and Healthy Control group (HC). ** p < 0.01; *** p < 0.001

### 3.3. Seed-based analyses

#### 3.3.1. Inferior frontal gyrus

At a group level, connectivity maps show that both groups have extended maps of fronto-temporal areas connected with the LIFG (Figure 2). They did not reveal any inter-group differences (threshold for second level maps p < 0.05 FWE at the cluster level). Regions positively and negatively correlated with the LIFG in each group are detailed in the Appendix.

**Figure 2.**
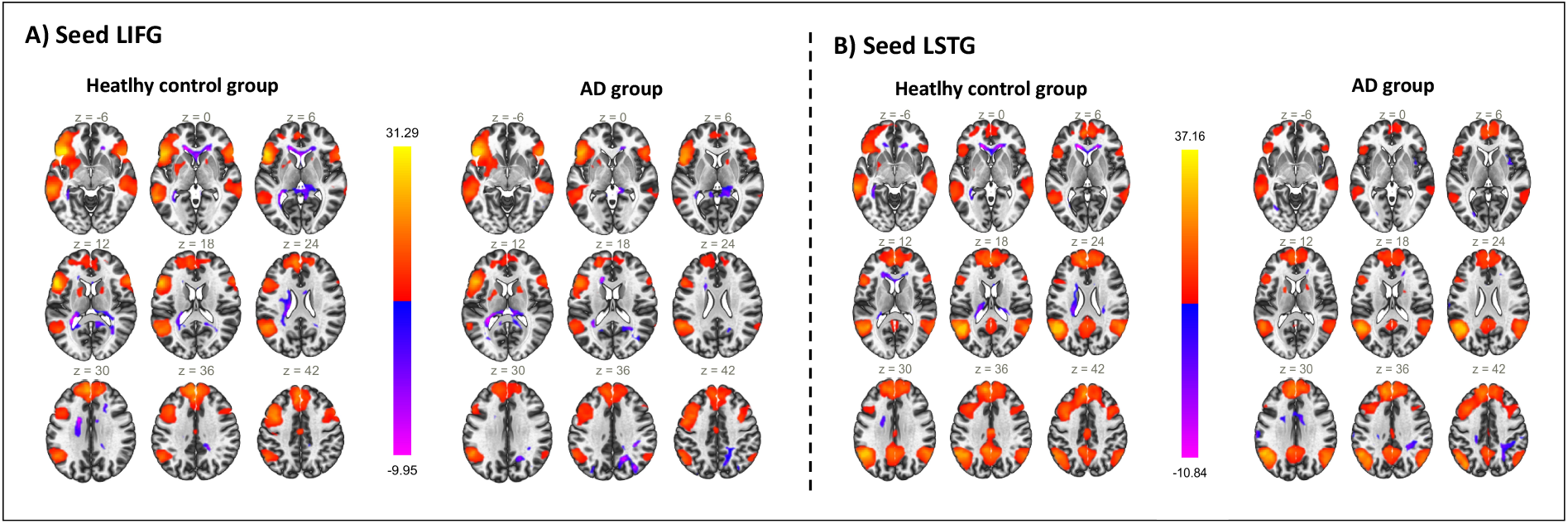
Cluster map for A) LIFG and B) LSTG in healthy controls and AD participants. Yellow to red color for clusters positively correlated to LIGF activity; blue to pink color for clusters negatively correlated to LIFG activity.

#### 3.3.2. Posterior temporal gyrus

Similar to the previous seed-based analysis, both groups had extended map areas connected with the LSTG (Figure 2). Two sample t-tests did not reveal any inter-group differences (threshold for second level maps p < 0.05 FWE at the cluster level). Regions positively and negatively correlated with the LIFG in each group are detailed in the Appendix.

### 3.4. Within- and between-network connectivity

The average connectivity within the language network (*t*(46)=1.12, *p=0*.*13*), within the Left ECN network (*t*(46)=1.35, *p=0*.*09*) and within the right ECN (*t*(46)=-0.77, *p=0*.*78*) was not significantly different between the two groups.

Additionally, the strength of connectivity between the language network and the left ECN network (*t*(46)=1., *p=0*.*46*) or between the language network and the right ECN network (*t*(46)=0.53, *p=0*.*3*) was not lower in the AD group.

### 3.5. Multivariate pattern analyses

#### 3.5.1. Language network

For the language structural network, the classification analysis yielded an accuracy of 95.8%. The permutation tests indicated that this classification was highly significant (p<0.0001). Furthermore, it indicated that the discriminative regions included the right inferior frontal gyrus, the right superior temporal gyrus, the left middle temporal gyrus and the left middle temporal gyrus/angular gyrus (Figure 3).

**Figure 3.**
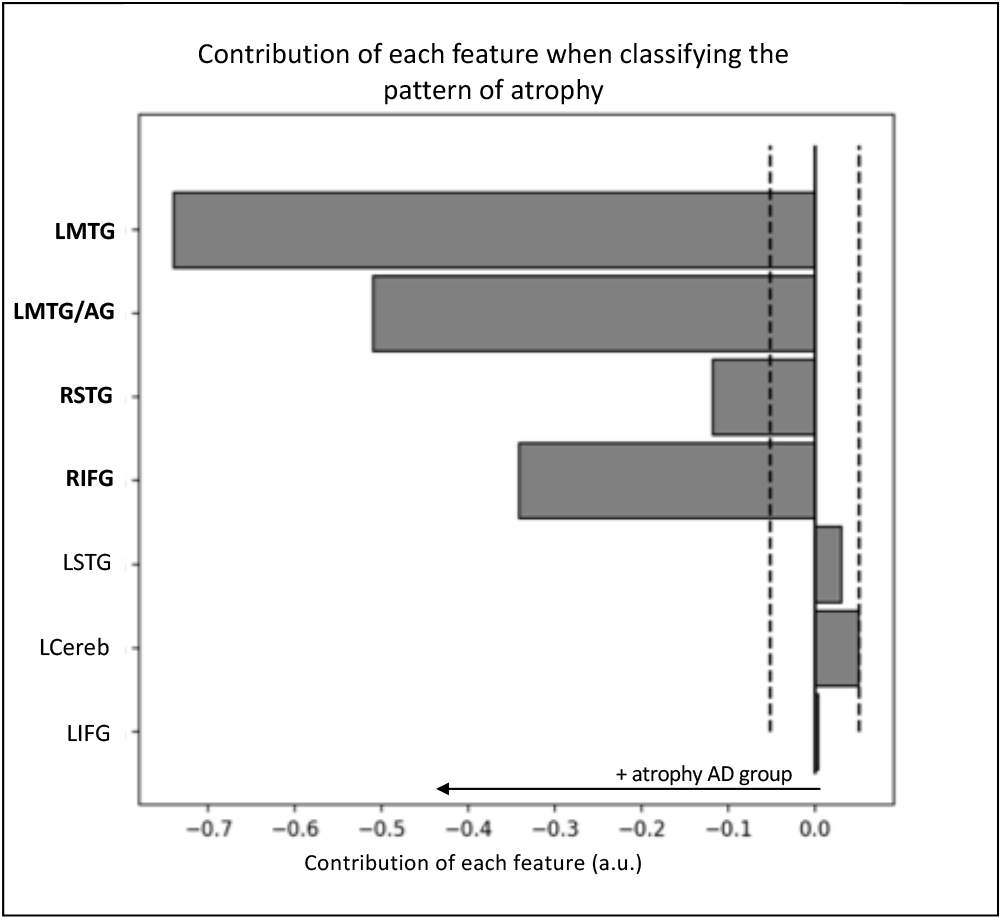
Contribution of each feature when classifying the pattern of atrophy within language network. The dashed lines represent the threshold (p<0.05) for significance obtained through permutations. Significant features are indicated in bold on the y-axis. Only the main area of each ROI is displayed on the y-axis. Abbreviations: LMTG: left middle temporal gyrus; LMTG/AG: left middle temporal/angular gyrus; LSTG: left middle/superior/supramarginal gyrus; RSTG: right middle/superior/ supramarginal gyrus, LCereb: left cerebellum.

Regarding the language functional network, the classification analysis yielded an accuracy of 64.5% (p<0.05). This pattern revealed a global increase in language functional connectivity in the AD group compared to the control group (Figure 4). There were three significantly discriminative connectivity features: the connectivity between the left inferior frontal gyrus and the left middle temporal gyrus/angular gyrus, the connectivity between the left inferior frontal gyrus and the left superior temporal gyrus and the connectivity between the left middle temporal gyrus/angular gyrus and the left superior temporal gyrus.

**Figure 4.**
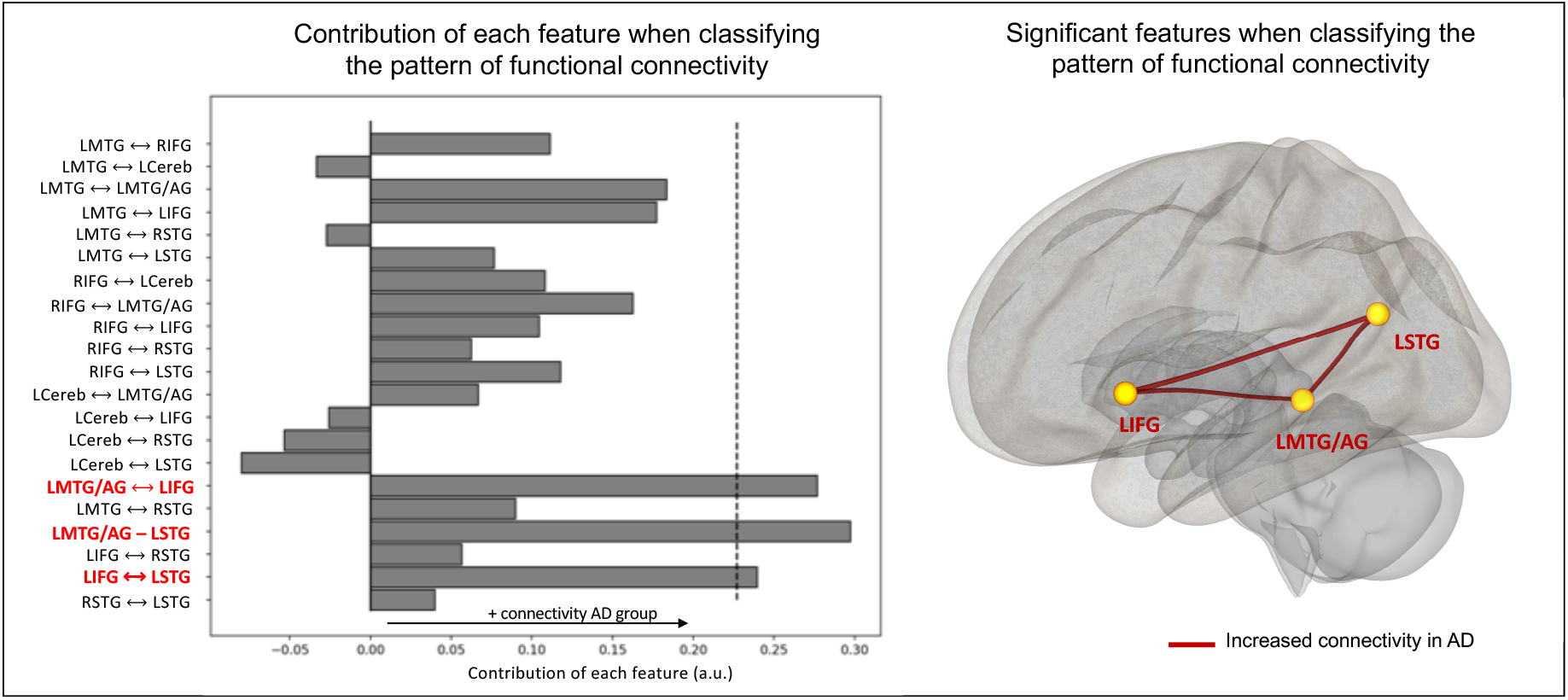
Contribution of each feature when classifying the pattern of functional connectivity within the language network. The dashed lines represent the threshold for significance (p<0.05) obtained through permutations. Significant features are indicated in bold red on the y-axis. Only the main area of each ROI is displayed on the y-axis. Abbreviations: LMTG: left middle temporal gyrus; LMTG/AG: left middle temporal/angular gyrus; LSTG: left middle/superior/supramarginal gyrus; RSTG: right middle/superior/ supramarginal gyrus, LCereb: left cerebellum.

#### 3.5.2. Executive control networks

For the structural left ECN, the classification analysis yielded an accuracy of 95.8% (p<0.0001). All the features of this network but one (left IFG/orbitofrontal gyrus) were significantly informative for the classification. For the structural right ECN, the classification analysis yielded an accuracy of 91.8% (p<0.0001). All the features of this network were significantly informative for the classification.

Regarding these functional networks, neither of the two networks could significantly discriminate AD participants from healthy controls (left ECN: 60.9%; p=0.09; right ECN: 44%; p=0.7).

#### 3.5.3. Correlations with language performance

We extracted a measure of group-typicality from classification analyses to perform intra-group correlations between discriminative patterns and language performance. For each participant, we extracted the confidence score of the classifier to predict the class of this participant (HC or AD). This score corresponds to the distance of each participant from the hyperplane that distinguishes the two classes. For instance, a participant whose data represent a point far from the classification hyperplane will have a high confidence score, indicating that they can be confidently classified as a member of the class (depending on which side of the hyperplane they fall). On the other hand, a participant whose data represent a point close to the classification hyperplane will have a low confidence score, indicating that this participant’s data were not very distinct from the other class. This measure therefore represents a continuous group-typicality measure that allow us to relate multivariate patterns analyses to behavioral performance (similarly to Ritchie and Carlson, 2016; Senoussi et al., 2016). For language performance, the most sensitive standardized language tasks (object naming, famous face naming, word spelling, written semantic verification, sentence spelling and text comprehension) and connected speech variables (lexical content, modalizing discourse, self-corrections) were chosen. Kendall correlations were performed, followed by Bonferroni-Holm corrections.

For the language structural network, confidence scores were not correlated with language performance, standardized tasks or the connected speech task in any group. For the language functional network, confidence scores were not correlated with any language task in the AD group. There was a positive correlation with the connected speech task in the HC group: participants with high confidence scores had superior lexical content during this task (p=0.015; r=0.36). This means that participants that were the most different from the AD group in terms of language functional connectivity had richer lexical content during their narrative production. Confidence scores obtained during structural ECN classifications were not correlated with language performance in any group.

## 4. Discussion

In the current study, we recruited typical AD participants at the prodromal stage who underwent a comprehensive language assessment, a structural 3D-T1 MRI and a resting-state fMRI. We showed that AD participants had language impairment during standardized language tasks and connected speech production. Based on MVPA results, an increased functional connectivity within the language network could be a marker of early AD, despite gray matter loss. However, such differences were not noticeable during univariate analyses.

### 4.1. Behavioral level

The prodromal AD group had lower performance than HC during several lexical tasks: object naming, famous face naming, word spelling and written semantic verification. These results are coherent with previous literature that showed an early semantic and naming impairment in AD (e.g. Barbeau et al., 2012; Joubert et al., 2010). Contrary to what was expected, their verbal fluency was not lower than in HC (contrary to Mueller et al., 2016).

During connected-speech production, the two groups did not differ in terms of number of words. Additionally, and similarly to Mueller et al. (2016), the prodromal AD group did not produce more filled pauses than HC. However, we revealed three qualitative differences in AD participants’ productions. First, their lexical content was lower than healthy controls, which is similar to what Pistono et al. (2019) found using the same narrative task. AD participants also produced more modalizing discourse and more self-corrections while speaking. Pistono et al. (2018) also found an increase of modalizing discourse in prodromal patients’ narratives. As mentioned by Duong et al. (2003), the fact that patients produced modalizing discourse means that their pragmatic abilities are preserved and used to communicate about their productions. It is therefore possible that this variable increases in prodromal AD but decreases in later stages, when pragmatic and metacognitive processes are altered. Similarly, self-corrections can be seen as evidence that some abilities remain. Indeed, self-corrections are the result of a relatively late process of verbal self-monitoring. Verbal self-monitoring is a cognitive system that inspects the speech plan and overt speech and initiates corrections when necessary (Hartsuiker, 2014). In the current study, the AD participants exhibited more errors than the controls. However, they were able to correct themselves, while an impaired monitoring system would lead to uncorrected errors. The significant proportion of modalizing discourse and self-corrections therefore reflects the use of metacognitive abilities in prodromal AD patients’ discourse production. In sum, in our sample, despite lexical difficulties, patients present with mostly preserved language/communicational abilities reflected by different compensation mechanisms during discourse production.

### 4.2. Univariate analyses

No inter-group differences were found during seed-based analyses, both when using the LIFG or LSTG as a seed. This result is, however, unsurprising, given the early stage of AD participants that were recruited in the current study. Indeed, Montembeault et al. (2019) recruited AD participants with a slightly lower MMSE than AD participants in the current study (24.9±3.1 in their study vs. 25.5±2.6 in the current study). They showed that only one cluster (the right posterior temporal gyrus) was significantly less connected to the left posterior temporal gyrus in prodromal AD, while there was no difference with the control group when the LIFG was used a seed.

More surprisingly, connectivity between the language network and the executive control network was not lower in AD participants. Since this type of measure is the result of mean pairwise correlations between several ROIs, it is possible that it is too broad to reveal differences at a prodromal stage. Additionally, the AD group did not significantly differ from the HC group on fluency tasks during language assessment (i.e. not after corrections for multiple comparisons). Studies that revealed interactions with the executive control network in healthy aging suggested that older adults may rely on these additional attentional resources to maintain successful verbal fluency performance (Muller et al., 2016; Pistono et al., 2020). It is therefore also possible that prodromal patients do not differ from HC in the interaction of language and executive resources. However, although the two groups did not differ on any of these univariate measures, the pattern of atrophy or functional connectivity within the language network helped discriminate the two groups, as shown with multivariate analyses.

### 4.3. Multivariate analyses

MVPA uses machine-learning algorithms that allow information patterns to be extracted from multi-dimensional data and the class of new data to be predicted. Here, we aimed to classify the two groups based on the pattern of atrophy and functional connectivity within the language network and the executive control networks. By doing so, we revealed two main findings. First, prodromal AD is not characterized by decreased language functional connectivity. Second, language network connectivity could better classify participants than executive control networks.

Regarding language networks, the pattern of atrophy was highly discriminative of AD participants from HC. However, this pattern was not correlated with language performance in any group. This discrepancy between atrophy and language performance has already been shown in the literature on healthy aging (Pistono et al., 2020). Additionally, while AD participants could be classified above chance based of their pattern of atrophy, the classifier was also able to discriminate them when examining their pattern of functional connectivity. However, this pattern revealed an overall increased connectivity between most language ROIs in the AD group. In other words, despite important gray matter loss, AD participants presented increased functional connectivity within language network. Taken individually, connectivity values between each ROI are not informative (i.e. not significantly different in univariate analyses); however, when all the information is considered, this global increase becomes discriminant. This pattern could have been caused by the fact that AD participants were at the prodromal stage. Indeed, increased functional connectivity associated with gray matter loss has already been shown in the literature about subjective cognitive impairment (Hafkemeijer et al., 2013) or mild cognitive impairment (Gardini et al., 2015). Two explanations are developed in the current literature: either this type of mechanism could compensate for cognitive decline, or increased functional connectivity reflects a shift in network properties that may cause further brain damage (Gallagher et al., 2010). This pattern of connectivity was not correlated with language performance in the AD group, while in HC, the confidence score of each individual was correlated with higher lexical content during connected-speech production. This means that HC that presented a pattern of connectivity highly different from AD participants had superior lexical content in their narrative. On the contrary, increased functional connectivity in prodromal AD does not seem sufficient to maintain behavioral performance. However, future work is required to examine whether increased connectivity switches to decreased connectivity at a later stage of AD and how it relates to language decline.

The pattern of atrophy in the left and right ECN was highly discriminative of AD participants from HC. However, similarly to the language network, this pattern was not correlated with language performance in any group. Additionally, classification accuracies of AD participants and HC based on the functional connectivity within executive control networks were not significant. This suggests that despite significant atrophy, AD participants’ functional connectivity patterns within the executive control networks were not different from HC.

Taken together, current findings show that language functional networks can better discriminate prodromal AD participants than executive control networks. More precisely, functional connectivity increased within AD participants’ language network, in particular between three areas: left IFG, left STG and left MTG/AG. While the language network is usually understudied in AD compared to other networks, it could provide important insight at an early stage.

### 4.4. Limitations

This study has 24 participants in each group, which is comparable to previous studies we mentioned earlier (e.g. Weiler et al., 2014), but represent a rather small sample size. Further studies are therefore required to examine structural and functional language network changes in prodromal AD and to reinforce current findings. Although we adapted our methods to the current sample size (e.g. using feature selection and cross-validations during MVPA), further research on large samples of participants could combine multiple modalities (e.g. language task performance, gray matter, functional connectivity) into a single multivariate pattern classification analysis. Moreover, as mentioned earlier, it would be interesting to replicate current methods on larger longitudinal data to uncover how the patterns we observed evolve over the course of AD.

Additionally, we did not use a functional language task to control that participants were left hemisphere dominant or to define our ROIs. Although we exclusively included right-handed participants, we cannot be sure that their language was left lateralized. Similarly, the use of a predefined atlas might have influenced the results. Nonetheless, we decided to use an atlas that was functionally defined, since these are more likely to represent brain regions effectively involved in language processing than anatomical seeds (Muller et al., 2014).

### 4.5. Conclusions

The current study demonstrated that prodromal participants present with language alterations, both when examining standardized language tasks and connected-speech production. It also showed that, when analyzing language functional networks, multivariate pattern analyses could significantly predict the group membership of prodromal patients and HC, while univariate analyses were not able to discriminate participants at this stage. This method therefore represents a useful tool for investigating the functional and structural (re-)organization of the neural bases of language in various populations.

## Data and code availability statement

Because of privacy issues regarding clinical data, neuroimaging data, raw language scores and discourse transcripts will be made available from the corresponding author upon reasonable request.

## Acknowledgments

This study was supported by a grant from the Occitania Region and the Toulouse Mind and Brain Institute to MJ (TellMA project grant number: 15050480). We would like to thank the Inserm/UPS UMR1214 Technical Platform for the MRI acquisitions. The authors would also like to thank the patients and control participants who participated in the study, as well as the promoter of the study, Toulouse University Hospital (CHU). There are no conflicts of interest to report.

## Appendix 1.

Regions showing less density of grey matter in AD participants in comparison to healthy controls. The statistical threshold is pFWE-corr<0.05 (k>50 voxels)

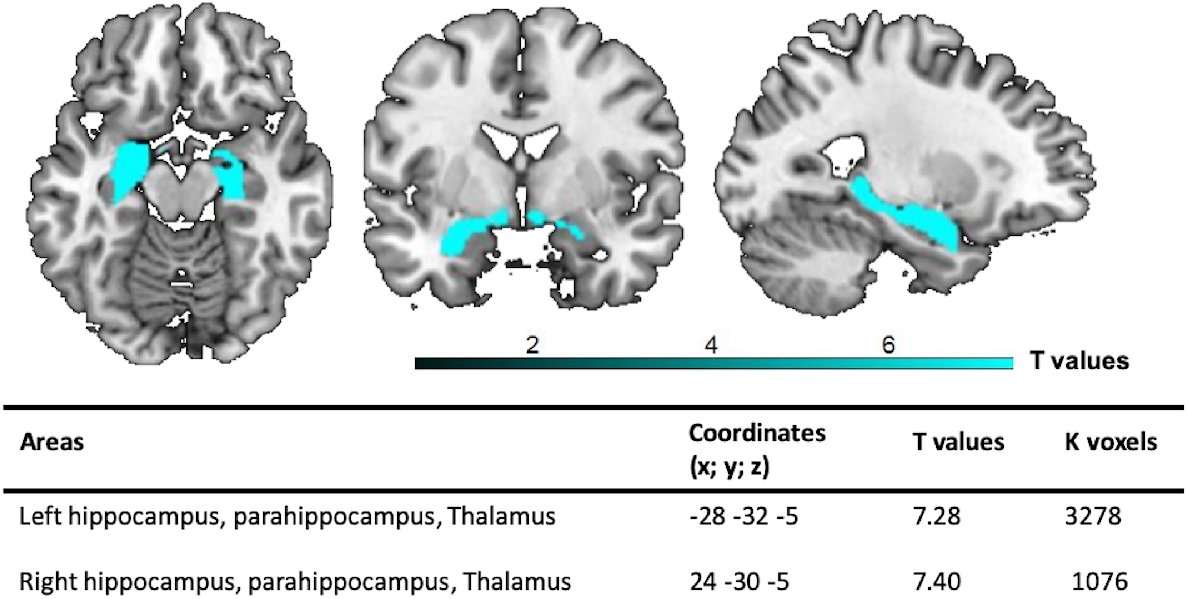

## Appendix 2.

Summary of regions positively and negatively correlated with the seeds in each group. Abbreviations: PCC = Posterior cingulate gyrus; SMG = Supramarginal gyrus; AG = Angular gyrus; TP = Temporal pole; ITG = Inferior temporal gyrus; MTG= Middle temporal gyrus; SPL= Superior parietal lobule.

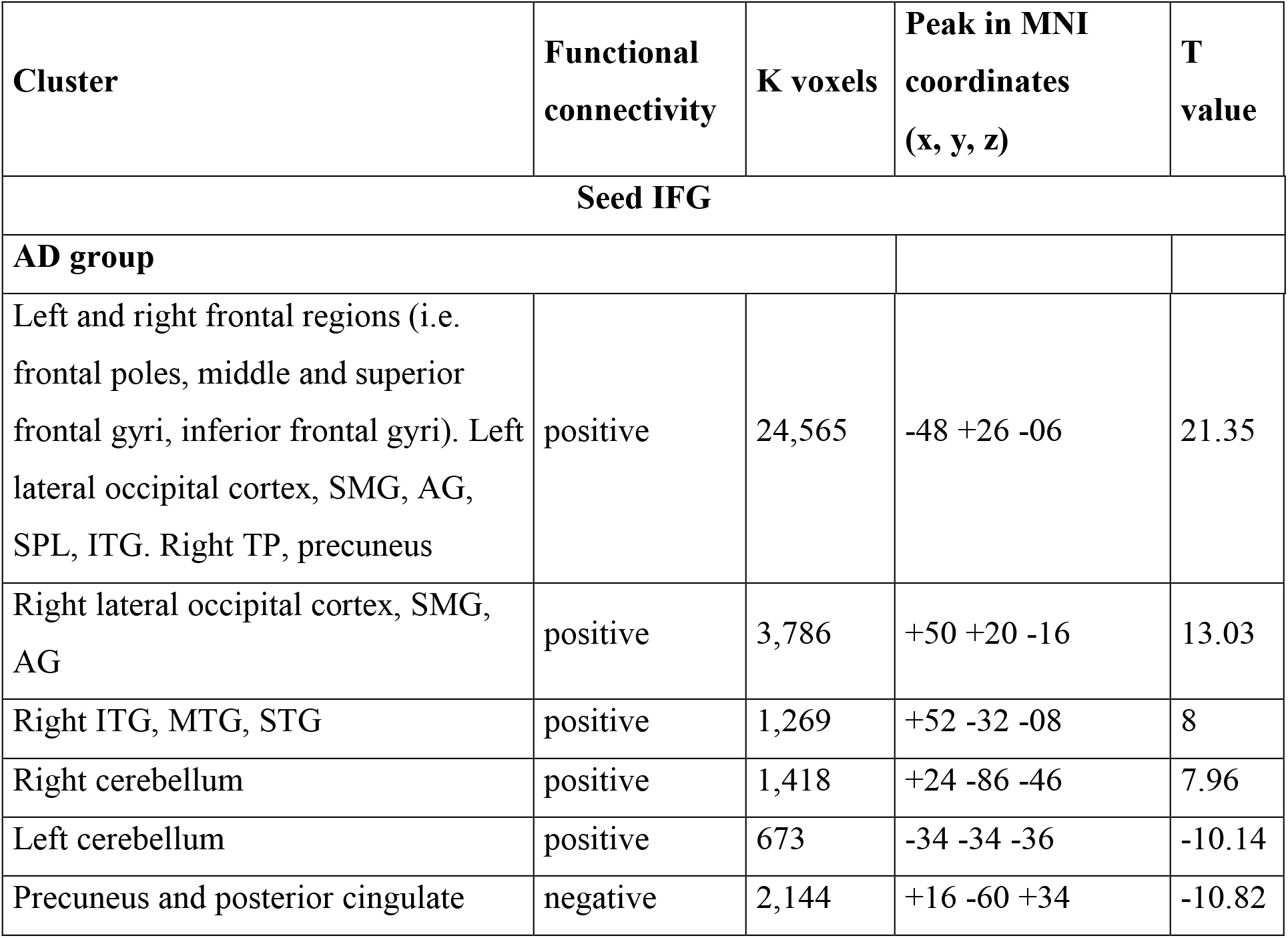
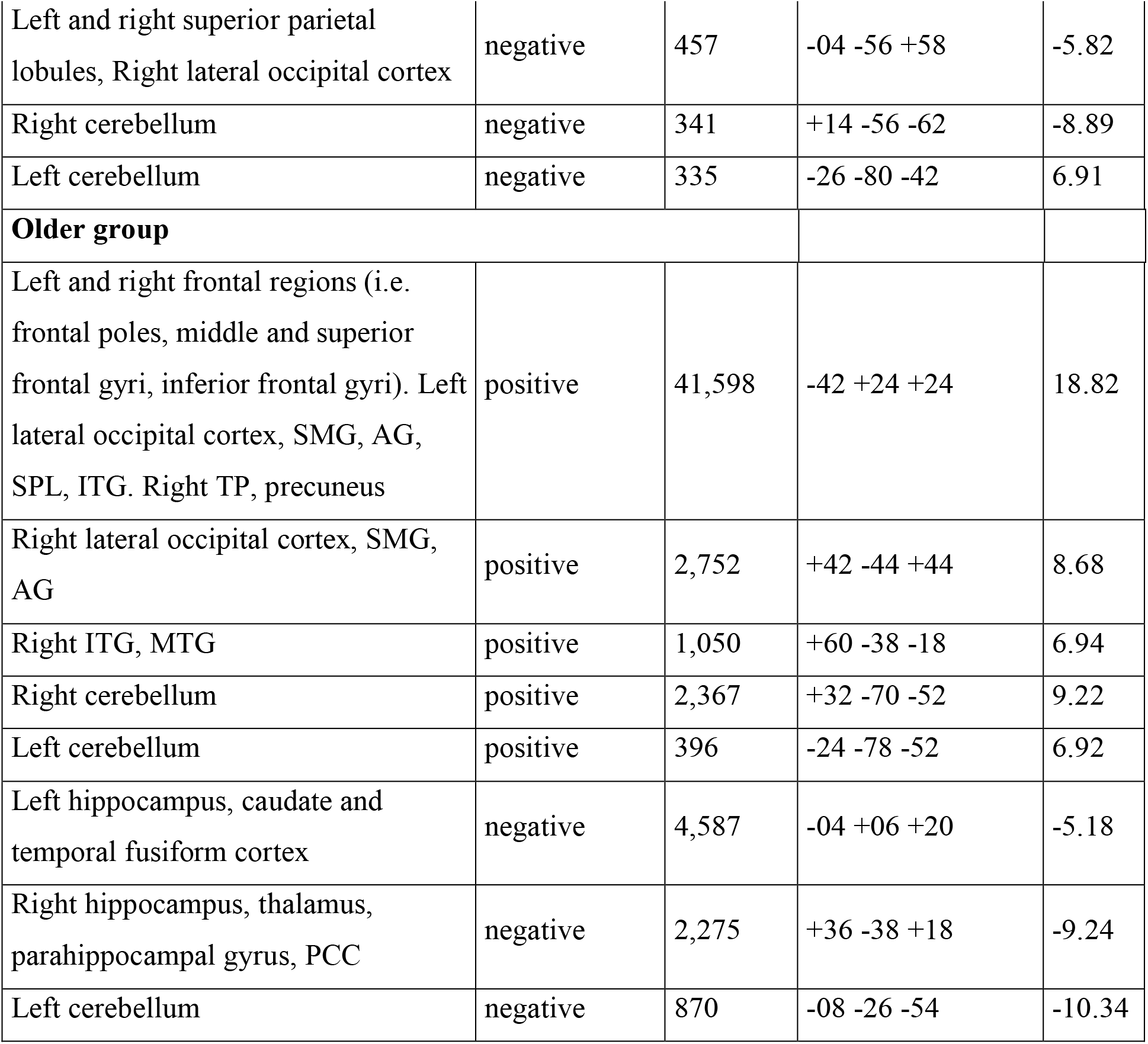

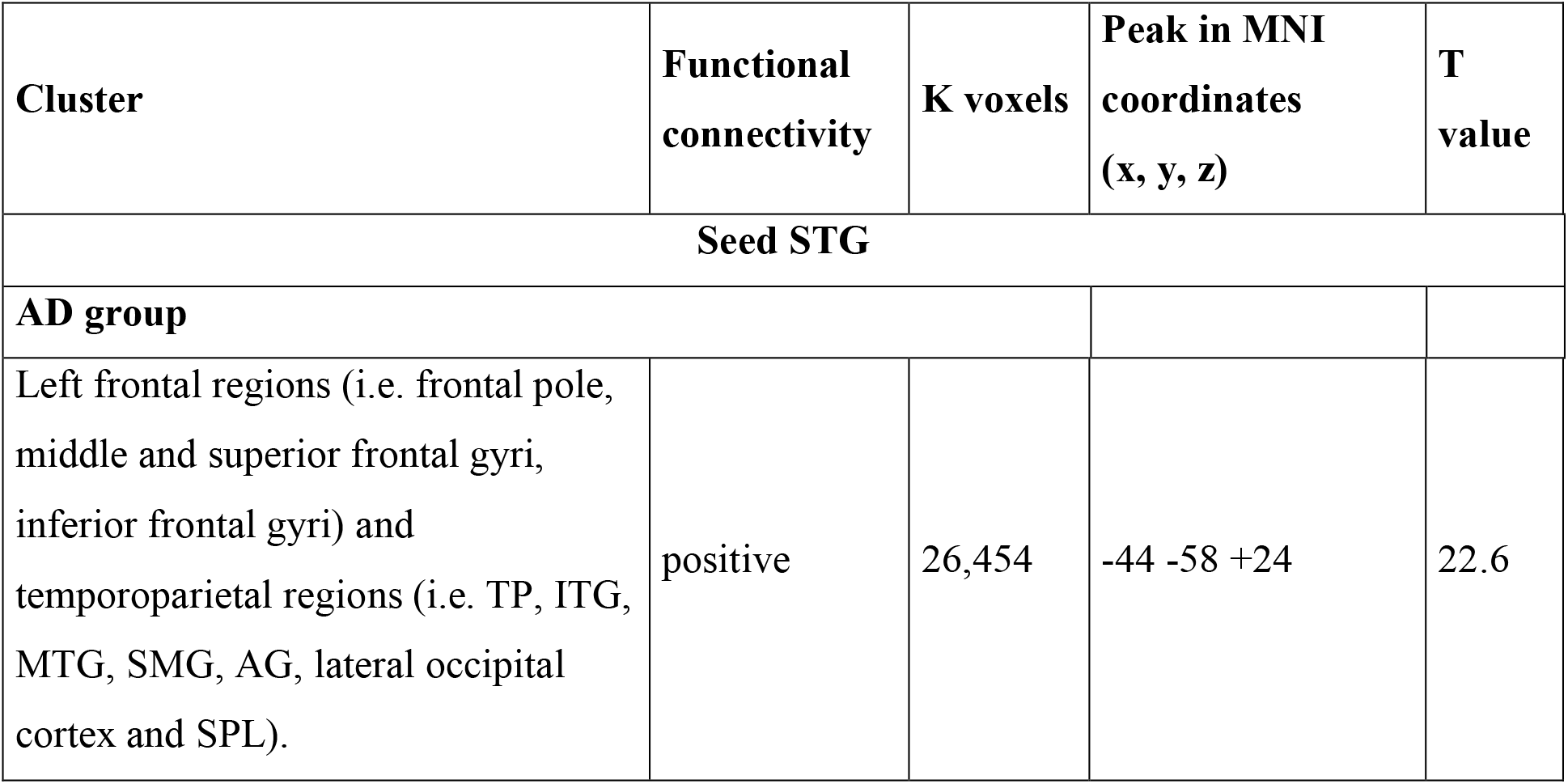
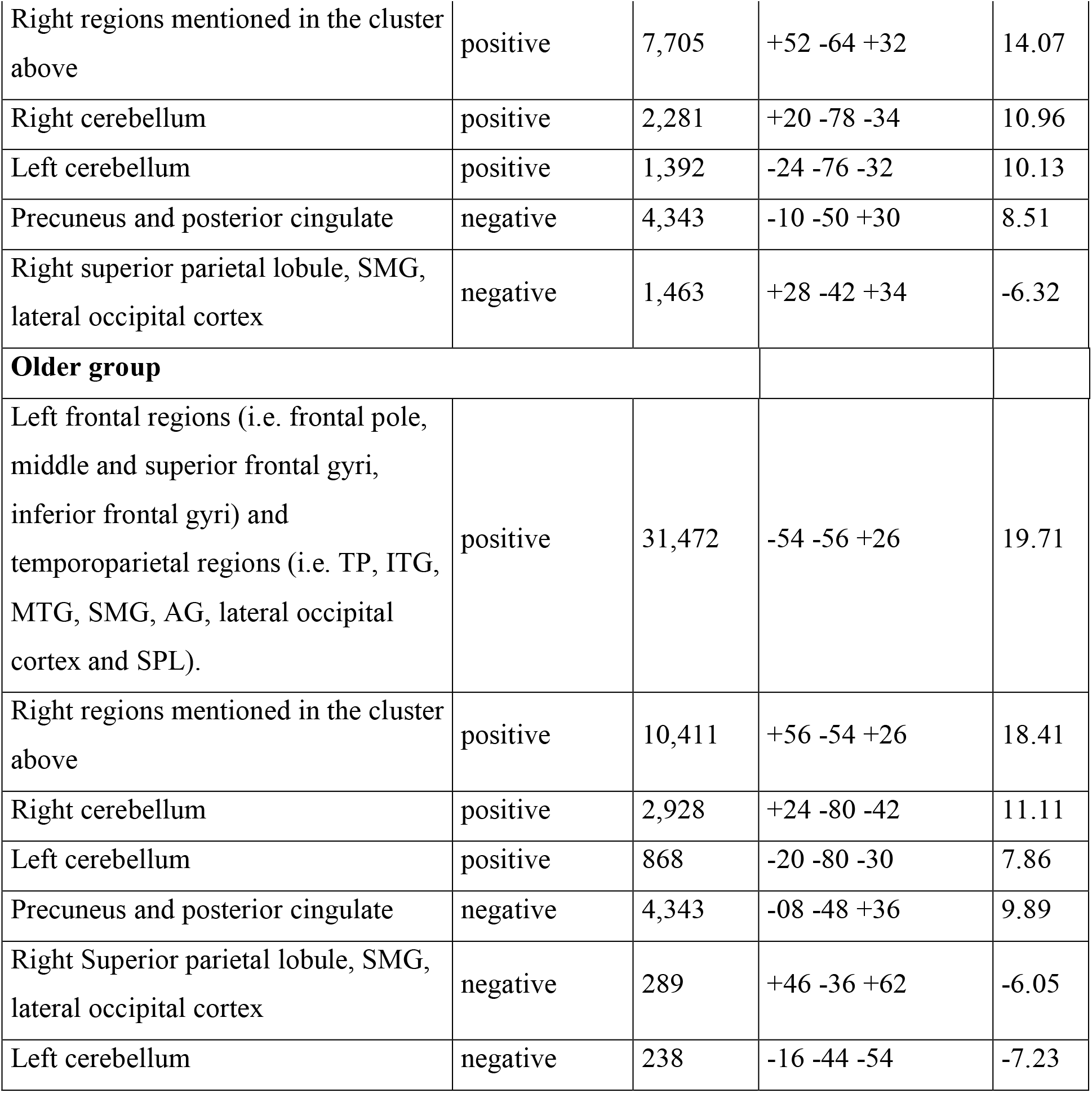

## References

Agniel, A., Joanette, Y., Doyon, B., Duchein, C., 1992. Protocole Montréal-Toulouse?: Évaluation des gnosies visuelles et auditives. Paris.

Ahmed, S., Haigh, A.F., Jager C.A. De, Garrard, P., 2013. Connected speech as a marker of disease progression in autopsy-proven Alzheimer’s disease. Brain. https://doi.org/10.1093/brain/awt269

Baddeley, A.D., Emslie, H., Nimmo-Smith, I., Company, T.V.T., 1994. Doors and People: A Test of Visual and Verbal Recall and Recognition. Manual. Thames Valley Test Company.

Barbeau, E.J., Didic, M., Joubert, S., Guedj, E., Koric, L., Felician, O., Ranjeva, J.P., Cozzone, P., Ceccaldi, M., 2012. Extent and neural basis of semantic memory impairment in mild cognitive impairment. J. Alzheimer’s Dis. 28, 823–837. https://doi.org/10.3233/JAD-2011-110989

Beck, A.T., Ward, C.H., Mendelson, M., Mock, J., Erbaugh, J., 1961. An inventory for measuring depression. Arch. Gen. Psychiatry 4, 561–71.

Bézy, C., Renard, A., Pariente, J., 2016. GREMOTS Batterie d’évaluation des troubles du langage dans les maladies neurodégénératives. De Boeck supérieur.

de Lira, J.O., Ortiz, K.Z., Campanha, A.C., Bertolucci, P.H.F., Minett, T.S.C., 2011. Microlinguistic aspects of the oral narrative in patients with Alzheimer’s disease. Int Psychogeriatr 23. https://doi.org/10.1017/S1041610210001092

Duong, A., Tardif, A., Ska, B., 2003. Discourse about discourse: What is it and how does it progress in Alzheimer’s disease? Brain Cogn. 53, 177–180. https://doi.org/10.1016/S0278-2626(03)00104-0

Gallagher, M., Bakker, A., Yassa, M.A., Stark, C.E.L., 2010. Bridging neurocognitive aging and disease modification: targeting functional mechanisms of memory impairment. Curr. Alzheimer Res. 7, 197–199. https://doi.org/10.2174/156720510791050867

Gardini, S., Venneri, A., Sambataro, F., Cuetos, F., Fasano, F., Marchi, M., Crisi, G., Caffarra, P., 2015. Increased functional connectivity in the default mode network in mild cognitive impairment: a maladaptive compensatory mechanism associated with poor semantic memory performance. J. Alzheimers. Dis. 45, 457–470. https://doi.org/10.3233/JAD-142547

Hafkemeijer, A., Altmann-Schneider, I., Oleksik, A.M., Van De Wiel, L., Middelkoop, H.A.M., Van Buchem, M.A., Van Der Grond, J., Rombouts, S.A.R.B., 2013. Increased functional connectivity and brain atrophy in elderly with subjective memory complaints. Brain Connect. 3, 353–362. https://doi.org/10.1089/brain.2013.0144

Hartsuiker, R., 2014. Monitoring and control of the production system. Oxford Handb. Lang. Prod. 417–436.

Haynes, J.-D., Rees, G., 2006. Decoding mental states from brain activity in humans. Nat. Rev. Neurosci. 7, 523–534. https://doi.org/10.1038/nrn1931

Hebart, M.N., Baker, C.I., 2018. Deconstructing multivariate decoding for the study of brain function. Neuroimage 180, 4–18. https://doi.org/10.1016/j.neuroimage.2017.08.005

Hoffman, P., Morcom, A.M., 2018. Age-related changes in the neural networks supporting semantic cognition: A meta-analysis of 47 functional neuroimaging studies. Neurosci. Biobehav. Rev. 84, 134–150. https://doi.org/10.1016/j.neubiorev.2017.11.010

Joubert, S., Brambati, S.M., Ansado, J., Barbeau, E.J., Felician, O., Didic, M., Lacombe, J., Goldstein, R., Chayer, C., Kergoat, M.-J., 2010. The cognitive and neural expression of semantic memory impairment in mild cognitive impairment and early Alzheimer’s disease. Neuropsychologia 48, 978–88. https://doi.org/10.1016/j.neuropsychologia.2009.11.019

Kemper, S., LaBarge, E., Farraro, R., Cheung, H., Storandt, M., 1993. On the preservation of syntax in Alzheimer’s disease. Arch. Neurol. 50, 81–86. https://doi.org/10.1001/archneur.1993.00540010075021

Liu, T., Wang, Y., Yan, T., 2018. Classification by a Rs-fMRI Study. 2018 11th Int. Congr. Image Signal Process. Biomed. Eng. Informatics 1–6.

Mahieux-Laurent, F., Fabre, C., Galbrun, E., Dubrulle, A., Moroni, C., 2008. Validation d’une batterie brève d’évaluation des praxies gestuelles pour consultation Mémoire. Évaluation chez 419 témoins, 127 patients atteints de troubles cognitifs légers et 320 patients atteints d’une démence. Rev. Neurol. (Paris). 1519, 511 YP–611. http://dx.doi.org/10.1016/j.neurol.2008.11.016

Mascali, D., Dinuzzo, M., Serra, L., Mangia, S., Maraviglia, B., 2018. Disruption of Semantic Network in Mild Alzheimer’s Disease Revealed by Resting-State fMRI. Neuroscience 371, 38–48. https://doi.org/10.1016/j.neuroscience.2017.11.030

Melrose, R.J., Campa, O.M., Harwood, D.G., Osato, S., Mandelkern, M.A., Sultzer, D.L., 2009. The neural correlates of naming and fluency deficits in Alzheimer’s disease: an FDG-PET study. Int. J. Geriatr. Psychiatry 24, 885–893. https://doi.org/10.1002/gps.2229

Montembeault, M., Chapleau, M., Jarret, J., Boukadi, M., Laforce, R., Wilson, M.A., Rouleau, I., Brambati, S.M., 2019. Differential language network functional connectivity alterations in Alzheimer’s disease and the semantic variant of primary progressive aphasia. Cortex 117, 284–298. https://doi.org/10.1016/j.cortex.2019.03.018

Mueller, K.D., Koscik, R.L., Turkstra, L.S., Riedeman, S.K., LaRue, A., Clark, L.R., Hermann, B., Sager, M.A., Johnson, S.C., 2016. Connected Language in Late Middle-Aged Adults at Risk for Alzheimer’s Disease. J. Alzheimer’s Dis. 54, 1539–1550. https://doi.org/10.3233/jad-160252

Muller, A.M., Mérillat, S., Jäncke, L., 2016. Older but still fluent? Insights from the intrinsically active baseline configuration of the aging brain using a data driven graph-theoretical approach. Neuroimage 127, 346–362. https://doi.org/10.1016/j.neuroimage.2015.12.027

Muller, A.M., Meyer, M., 2014. Language in the brain at rest: new insights from resting state data and graph theoretical analysis.Front. Hum. Neurosci. 8, 228. https://doi.org/10.3389/fnhum.2014.00228

Pedregosa, F., Varoquaux, G., Gramfort, A., Michel, V., Thirion, B., Grisel, O., Blondel, M., Prettenhofer, P., Weiss, R., Dubourg, V., et al., 2011. Scikit-Learn: Machine Learning in Python. J. Mach. Learn. Res. 12, 2825–2830.

Pereira, F., Mitchell, T., Botvinick, M., 2009. Machine learning classifiers and fMRI: a tutorial overview. Neuroimage 45. https://doi.org/10.1016/j.neuroimage.2008.11.007

Pistono, A., Guerrier, L., Péran, P., Rafiq, M., Gimeno, M., Bézy, C., Pariente, J., Jucla, M., 2020. Increased functional connectivity supports language performance in healthy aging despite grey matter loss. Neurobiol. Aging. https://doi.org/10.1016/j.neurobiolaging.2020.09.015

Pistono, A., Jucla, M., Pariente, J., 2018. Discourse macrolinguistic impairment as a marker of linguistic and extralinguistic functions decline in early Alzheimer’s disease, International Journal of Language & Communication Disorders. https://doi.org/10.1111/1460-6984.12444

Pistono, A., Pariente, J., Bézy, C., Lemesle, B., Le Men, J., Jucla, M., 2019. What happens when nothing happens? An investigation of pauses as a compensatory mechanism in early Alzheimer’s disease. Neuropsychologia 124, 133–143. https://doi.org/10.1016/J.NEUROPSYCHOLOGIA.2018.12.018

Reitan, R.M., 1958. Validity of the Trail Making Test as an indicator of organic brain damage. Percept. Mot. Skills 8, 271–276. https://doi.org/10.2466/PMS.8.7.271-276

Ritchie, J.B., Carlson, T.A., 2016. Neural Decoding and “Inner” Psychophysics: A Distance-to-Bound Approach for Linking Mind, Brain, and Behavior. Front. Neurosci. 10, 190. https://doi.org/10.3389/fnins.2016.00190

Senoussi, M., Berry, I., VanRullen, R., Reddy, L., 2016. Multivoxel Object Representations in Adult Human Visual Cortex Are Flexible: An Associative Learning Study. J. Cogn. Neurosci. 28, 852–868. https://doi.org/10.1162/jocn_a_00933

Shirer, W.R., Ryali, S., Rykhlevskaia, E., Menon, V., Greicius, M.D., 2012. Decoding subject-driven cognitive states with whole-brain connectivity patterns. Cereb. Cortex 22, 158–165. https://doi.org/10.1093/cercor/bhr099

Starkstein, S.E., Mayberg, H.S., Preziosi, T.J., Andrezejewski, P., Leiguarda, R., Robinson, R.G., 1992. Reliability, validity, and clinical correlates of apathy in Parkinson’s disease. J. Neuropsychiatry Clin. Neurosci. 4, 134–9. https://doi.org/10.1176/jnp.4.2.134

Taler, V., Phillips, N.A., 2008. Language performance in Alzheimer’s disease and mild cognitive impairment: a comparative review. J. Clin. Exp. Neuropsychol. 30, 501–556. https://doi.org/10.1080/13803390701550128

Wechsler, D., 1997. WAIS-III: Administration and Scoring Manual. The Psychological Corporation, San Antonio:

Weiler, M., Fukuda, A., Massabki, L., Lopes, T., Franco, A., Damasceno, B., Cendes, F., Balthazar, M., 2014. Default Mode, Executive Function, and Language Functional Connectivity Networks are Compromised in Mild Alzheimer’s Disease. Curr. Alzheimer Res. 11, 274–282. https://doi.org/10.2174/1567205011666140131114716

Whitfield-Gabrieli, S., Nieto-Castanon, A., 2012. Conn: A Functional Connectivity Toolbox for Correlated and Anticorrelated Brain Networks. Brain Connect. 2, 125–141. https://doi.org/10.1089/brain.2012.0073

